# Aging redirects prior expectations from perception to action

**DOI:** 10.64898/2026.01.09.698574

**Authors:** Luca Tarasi, Margherita Covelli, Chiara Tabarelli de Fatis, Caterina Bertini, Alessio Avenanti, Vincenzo Romei

**Author notes:** **Corresponding Author:** Vincenzo Romei, Centro Studi e Ricerche in Neuroscienze Cognitive, Dipartimento di Psicologia, Alma Mater Studiorum – Università di Bologna, Campus di Cesena, via Rasi e Spinelli, 176. 47521 Cesena, Italy., Luca Tarasi, Centro Studi e Ricerche in Neuroscienze Cognitive, Dipartimento di Psicologia, Alma Mater Studiorum – Università di Bologna, Campus di Cesena, via Rasi e Spinelli, 176. 47521 Cesena, Italy.

## Abstract

Aging profoundly reshapes how the brain integrates sensory evidence with prior expectations during perceptual decision-making. Combining behavioural modelling with high-density EEG, we show that older adults assign greater weight on prior information, resulting in stronger decisional bias and reduced sensory precision under probabilistic cues. Neural dynamics revealed a breakdown of anticipatory tuning in visual alpha rhythms, supporting sensory preparation in youth, alongside enhanced central beta activity, indicating a shift toward pre-emptive motor engagement aligned with expected responses. A data-driven decomposition further showed that, whereas prior-driven behaviour in younger adults emerges from a flexible interplay between sensory and executive oscillatory systems, it becomes dominated in aging by a rigid, response-centred mode. Together, these findings uncover a large-scale reconfiguration of predictive computations with age, marking a transition from adaptive sensory-executive coordination to a more automatized, action-oriented predictive strategy.

## Introduction

Perceptual decision-making relies on the brain’s ability to anticipate sensory input based on prior experience, integrating top-down expectations with incoming evidence^1^. In young adults, extensive evidence shows that prior expectations systematically bias perception toward predicted sensory events, optimizing performance under uncertainty by assigning greater weight to expected than to unexpected inputs^2^. For example, probabilistic cues indicating the likely presence of a target tilt both perceptual decisions and confidence toward the cued alternative, without increasing overall accuracy^3–5^. These findings indicate that prior information shapes not only perceptual but also metacognitive judgments, engaging mechanisms partially dissociable from sensory discrimination itself.

How this inferential balance changes with age remains unclear. We hypothesize that aging biases perceptual inference toward a prior-weighted policy. As sensory sampling degrades with age, external evidence is underweighted, allowing expectations to exert greater influence and increasing prior-congruent responding. We further predict that this behavioural shift is mirrored by age-related changes in the neural computations that implement predictive inference.

Converging frameworks identify coordinated oscillatory dynamics across sensory, motor and executive systems as core substrates of predictive processing: posterior alpha supports perceptual preparation for expected inputs and sensory sampling; sensorimotor beta sustains and pre-activates action sets aligned with predictions; and fronto-parietal theta provides executive monitoring that avoid prior overweighting. Within the sensory domain, posterior alpha oscillations act as gatekeepers, dynamically regulating sensory cortical excitability according to stimulus expectations^6–11^. For instance, when cues indicate high versus low target probability, alpha amplitude scales with expected likelihood, predicting the extent to which priors bias perceptual decisions. Beyond amplitude, the individual frequency of alpha activity (IAF) indexes the temporal resolution of sensory sampling^12,13^. This rhythm typically slows with age^14,15^, potentially reducing the precision of perceptual encoding^16^ and thereby promoting stronger reliance on prior expectations. In parallel, expectations also engage the motor system by pre-activating response plans aligned with predicted outcomes^17–20^. This preparatory state appears as pre-stimulus beta desynchronization over motor regions when priors specify the most probable action^21^. By contrast, fronto-parietal theta supports executive monitoring, constraining over-reliance on priors to preserve flexible, context-sensitive decisions^11,22,23^.

Building on this framework, we investigated whether aging shifts inference from distributed, flexible integration to a more deterministic, action-oriented mode. We tested a group of younger (N = 80) and older (N = 72) adults performed a probabilistic detection task in which a pre-stimulus cue signalled target likelihood while EEG recorded the neural implementation of prior expectations.

We predicted (1) stronger prior-driven response biases, reflecting an over-weighting of internal expectations, and (2) reduced perceptual accuracy, indicating that this bias comes at the expense of sensory evidence processing. At the neural level, we expected (3) attenuated modulation of posterior alpha oscillations, indexing less efficient perceptual preparation; 4) a global slowing of the IAF, reflecting reduced precision of sensory sampling; (5) enhanced motor-beta desynchronization, reflecting a greater tendency to pre-activate motor responses aligned with prior expectations; and (6) altered fronto-parietal theta connectivity, indicating changes in executive coordination during the integration of priors.

## Results

### Older adults show reduced sensitivity and increased bias in the presence of prior expectations

Behavioural analyses were performed using Signal Detection Theory (SDT) to quantify perceptual sensitivity (d’) and response criterion (bias). We aimed to understand how aging affects the ability to integrate prior expectations into perceptual decisions. Two-way mixed ANOVAs (Group: young, older adults; Cue Probability: low, medium, high) showed significant interactions for both criterion (F _2,300_ = 8.29, p < 0.001) and sensitivity (F _2,300_ = 3.41, p = 0.034) measures. The analysis revealed that both age groups showed cue-congruent modulations in their criterion. Specifically, participants adopted a significantly more conservative decision bias (i.e., decreased tendency in reporting target presence) following low-probability cues (young = 0.63 ± 0.05; elderly = 0.67 ± 0.05), and a significantly more liberal bias (i.e., increased tendency to report target presence) following high-probability cues (young = -0.20 ± 0.05; old = -0.45 ± 0.09) compared to the medium probability condition (young = 0.37 ± 0.05, all t79 > 7.15, all p < 0.001; old = 0.10 ± 0.05, all t71 > 6.13, all p < 0.001). Additionally, between-group comparisons indicated that older adults adopted a significantly more liberal decision bias under medium and high probability conditions relative to young adults (all t_150_ > 4.16, all p < 0.001). To quantify the magnitude of this strategic shift, we computed a Δ*criterion* index reflecting the change in response bias between low- and high-probability cues. This analysis revealed a significantly larger Δ*criterion* in older adults than in young adults (young = 0.65 ± 0.07; elderly = 1.12 ± 0.15, t₁₅₀ = 2.89, p < 0.01]), confirming that aging is associated with an enhanced modulation of decision strategy by prior information. Age-related differences were also found when considering perceptual sensitivity. Specifically, older adults exhibited significantly reduced d’ in the high-probability condition (1.11 ± 0.10) compared to both the medium (1.34 ± 0.10) and low (1.26 ± 0.11) probability conditions (all t71 > 2.80, all p < 0.007), indicating that greater reliance on prior expectations negatively impacted perceptual accuracy in older individuals. In contrast, no significant cue-related modulation of d’ was observed in young adults (low = 1.36 ± 0.07; medium = 1.41 ± 0.07; high = 1.34 ± 0.10; all t_79_ < 1.53, all p > 0.133), nor between the two groups (all t_150_ < 1.87, all p > 0.064). Importantly, despite showing reduced perceptual sensitivity under high-probability conditions, older adults still performed the task well-above chance level. A one-sample t-test confirmed that d′ was significantly greater than zero (t₇₁ > 11.08, p < 0.001), indicating preserved basic perceptual discriminability.

**Figure 1.**
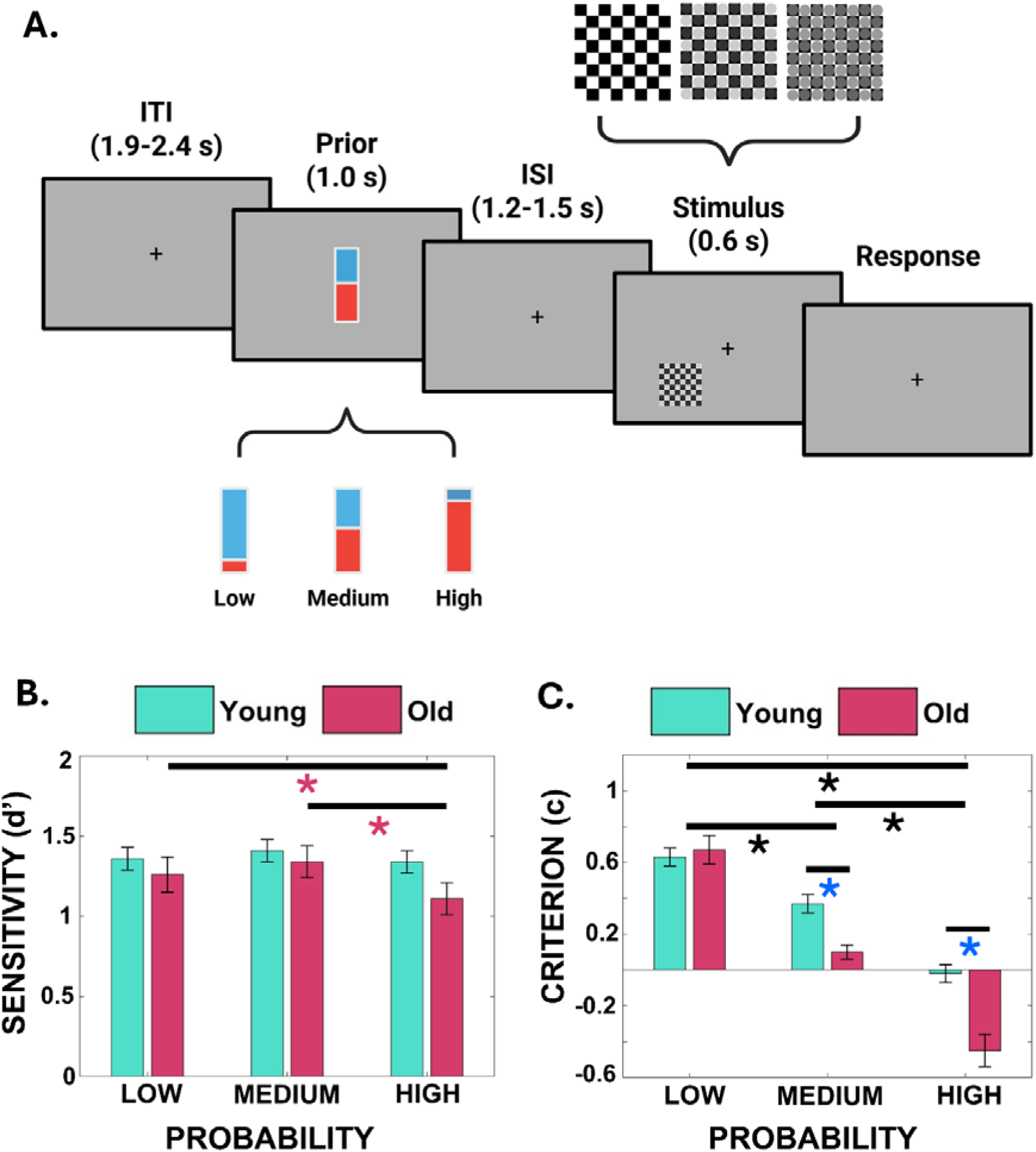
Task design and behavioral results. A. Participants completed a probabilistic detection task. Each trial began with the presentation of a probabilistic cue (low = 33 %, medium = 50 %, high = 67 %), followed 1.2-1.5 s later by the checkerboard stimulus in the lower-left quadrant containing either the target (grey circles) or a catch trial. Participants reported presence or absence of the target with their right hand. B. Signal-Detection Theory analyses revealed significant Group × Cue Probability interactions for sensitivity (F_₂_,_₃₀₀_ = 3.41, p = 0.034). High probability cues impaired d′ in older adults (purple asterisks, high 1.11 ± 0.10 vs. medium 1.34 ± 0.10 and low 1.26 ± 0.11; all t_₇₁_ > 2.80, p < 0.007), whereas young adults’ d′ remained stable across cues (all p > 0.133). No overall group difference in d′ emerged (young 1.37 ± 0.07 vs. old 1.24 ± 0.10; F_₁_,_₁₅₀_ = 1.33, p = 0.251). C. Signal-Detection Theory analyses revealed significant Group × Cue Probability interactions for decision criterion (F_₂_,_₃₀₀_ = 8.29, p < 0.001). A main effect of cue (black asterisks) demonstrated that both young and older adults adopted more conservative biases after low-probability cues (young c = 0.63 ± 0.05; old = 0.67 ± 0.05) and more liberal biases after high-probability cues (young c = –0.20 ± 0.05; old = –0.45 ± 0.09) versus medium (all t > 6.13, p < 0.001). Older adults exhibited significantly more liberal biases than young in medium/high conditions (blue asterisk, all t_₁₅₀_ > 4.16, p < 0.001).

### Alpha amplitude encodes probabilistic cues in young but not older adults

We next examined four oscillatory markers indexing distinct stages of predictive processing: posterior alpha amplitude, reflecting sensory anticipation; individual alpha frequency (IAF), indexing intrinsic sensory sampling; motor beta desynchronization, linked to anticipatory motor engagement; and fronto-parietal theta connectivity, associated with executive monitoring ^18,21^.

For posterior alpha amplitude, we focused on pre-stimulus alpha amplitude over a right posterior ROI (Oz, POz, Pz, O2, PO4, PO8, P2, P4, P6, P8), consistent with the contralateral engagement of visual cortex given that stimuli were presented in the left hemifield. A two-way mixed ANOVA with Group (young, older adults) as the between-subject factor and Cue Probability (low, medium, high) as the within-subject factor revealed a significant Group × Cue interaction (*F*₂,₃₀₀ = 4.65, *p* = 0.010) and a main effect of Cue (*F*₂,₃₀₀ = 3.63, *p* = 0.028), whereas the main effect of Group was not significant (*F*₁,₁₅₀ = 1.77, *p* = 0.185). Follow-up analyses showed that this effect was driven by significant alpha modulation in young adults. Specifically, this group was characterized by a marked reduction in posterior alpha amplitude under high-probability cues (–0.63 ± 0.06), compared to both low (–0.53 ± 0.06) and medium (–0.54 ± 0.06) conditions (all t79 > 3.32, all p < 0.001), which did not differ significantly from each other (t79 = 0.48, p = 0.63). This pattern reflects effective sensory preparation in response to prior expectations signalling target presence. In contrast, older adults showed no significant differences in alpha amplitude across cue levels (low = –0.46 ± 0.06; medium = –0.45 ± 0.06; high = –0.45 ± 0.05; all t₇₁ < 0.354, p > 0.722), suggesting attenuated modulation in their sensory activity by probabilistic information. Importantly, a between-group comparison demonstrated a significant difference in the high-probability condition (t₁₅₀ = 2.12, p = 0.036), with young adults showing stronger alpha suppression than older adults. Together, these findings show that age-related differences in anticipatory alpha suppression emerge specifically under high-probability contexts, indicating that older adults fail to flexibly tune sensory preparatory activity to the predictive structure of the environment. This impaired adjustment may underlie their reduced perceptual readiness and a compensatory shift toward downstream, motor-based strategies.

**Figure 2.**
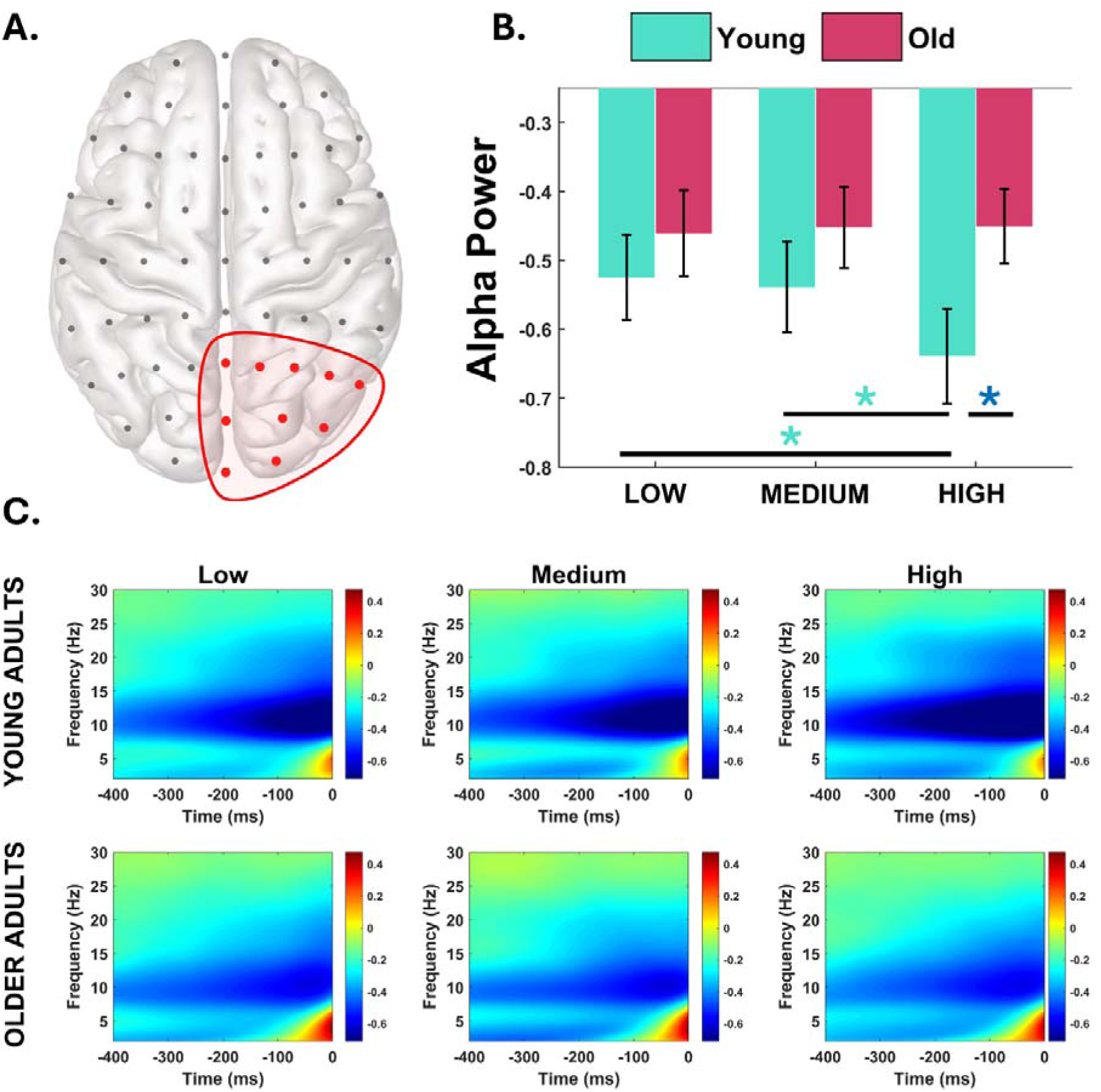
Posterior alpha___band modulation reflects prior___induced decision bias in young but not older adults. A. Pre-stimulus alpha amplitude (7–13 Hz) was extracted from posterior electrodes, corresponding to the hemisphere contralateral to stimulus presentation (Oz, POz, Pz, O2, PO4, PO8, P2, P4, P6, P8) for low (33 %), medium (50 %), and high (67 %) cue-probability conditions, separately for young and older adults. B. A significant Group × Cue Probability interaction (F_₂_,_₃₀₀_ = 4.35, p = 0.014) was driven by a pronounced alpha reduction under high-probability cues in young adults (green asterisks; high = –0.67 ± 0.07 vs. low = –0.55 ± 0.07 and medium = –0.57 ± 0.07; all t_₇₉_ > 3.11, p < 0.003), whereas older adults showed no cue-dependent modulation (all t_₇₁_ < 0.28, p > 0.78). In the high-probability condition, young adults also exhibited significantly stronger alpha desynchronization than older adults (blue asterisk, t_₁₅₀_ = 2.12, p = 0.036). C. Time–frequency representation of pre-stimulus activity (–400 to 0 ms) over posterior electrodes. Prior information enhanced sensory preparation through cue-dependent modulation of alpha amplitude, an effect markedly reduced in older adults, whose alpha power remained stable across cues, indicating attenuated sensory-preparatory dynamics during expectation-guided perception.

### Individual Alpha Frequency (IAF) is reduced in older adults across all cue conditions

The analyses of the individual alpha frequency (IAF) examined whether probabilistic cues modulated the intrinsic oscillatory peak across age groups. The two-way mixed ANOVA with factors Group (young, older adults) and Cue Probability (low, medium, high) revealed a significant main effect of Group (F₁,₁₅₀ = 6.06, p = 0.015), with older adults exhibiting a slower alpha rhythm (10.39 ± 0.09 Hz) than younger participants (10.79 ± 0.07 Hz). Neither the main effect of Cue nor the Group × Cue interaction reached significance (all F₂,₃₀₀ < 2.73, all p > 0.067), indicating no reliable cue-dependent modulation in either group. These results demonstrate that aging is associated with a global slowing of intrinsic alpha frequency, consistent with reduced efficiency of the sensory sampling process. Such slowing may reflect a coarser temporal resolution in integrating sensory evidence, providing a neural basis for the shift toward more expectation-driven perceptual strategies observed behaviourally.

**Figure 3.**
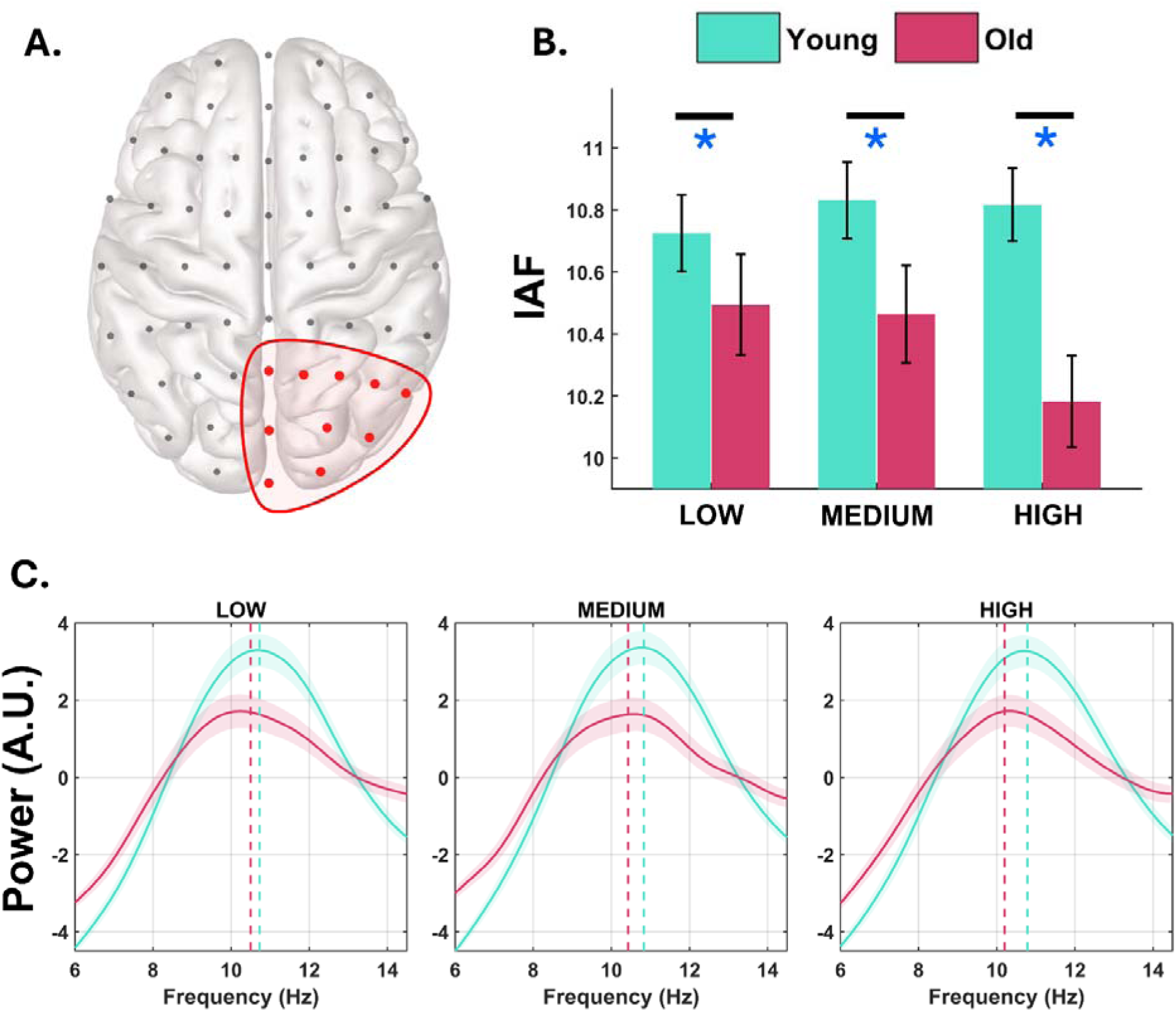
Individual alpha frequency by age group and cue probability. A. Individual alpha frequency (Hz) was extracted from posterior electrodes (Oz, POz, Pz, O2, PO4, PO8, P2, P4, P6, P8) for low (33 %), medium (50 %), and high (67 %) cue-probability conditions, separately for young and older adults as in the alpha amplitude analysis. B. A two-way mixed ANOVA revealed a significant main effect of Group (F_₁_,_₁₅₀_ = 6.06, p = 0.015), indicating generally lower IAF values in older compared with younger adults (blue asterisks, 10.39 ± 0.09 vs. 10.79 ± 0.07 Hz). Neither the main effect of cue probability nor the Group × Cue interaction reached significance (all F_₂_,_₃₀₀_ < 2.73, all p > 0.067), suggesting the absence of cue-dependent modulation in either group. C. Power spectrum within the alpha band across groups. Aging is associated with a global slowing of the individual alpha frequency (IAF), reflected in a downward shift of the spectral peak toward lower frequencies, consistent with reduced intrinsic sampling speed in the older cohort.

### Age-related increase in beta desynchronization in response to probabilistic cues

While posterior alpha suppression indexed efficient sensory preparation in young adults, its absence in older adults suggests a shift away from sensory-driven strategies. This raises the possibility that aging may involve a compensatory reliance on downstream systems, particularly within the motor domain. To test this possibility, we examined pre-stimulus beta amplitude over a left-central ROI (C5, C3, C1, CP5, CP3, CP1), contralateral to the responding right hand, to assess whether prior information modulated preparatory motor dynamics. A two-way mixed ANOVA revealed a significant main effect of cue probability (*F*₂,₃₀₀ = 17.32, *p* < 0.001) and a significant main effect of group (*F*₁,₁₅₀ = 12.62, *p* < 0.001), while the Group × Cue interaction was not significant (*F*₂,₃₀₀ = 0.70, *p* = 0.498). Across groups, beta desynchronization was significantly stronger in the informative conditions (high = –0.37 ± 0.04; low = –0.35 ± 0.04) compared to the medium probability condition (–0.29 ± 0.04; all t_151_ > 4.07, all p < 0.001), with no significant difference between the informative cues (t151 = 1.297, all p = 0.196). However, exploratory analysis showed a trend toward stronger beta desynchronization in the high-probability condition relative to the low-probability in older adults only (t71 = -1.778, p = 0.080). This pattern suggests that the mere presence of prior information enhances motor preparation, likely through the anticipatory activation of motor plans. In contrast, no such preparation is possible in the medium probability condition, where no directional bias can be formed. Crucially, the main effect of Group was driven by a general enhancement of beta suppression in older adults across all conditions relative to younger participants (young = –0.20 ± 0.03; old = –0.44 ± 0.03). Together, these findings indicate that aging amplifies motor preparatory engagement, as reflected by stronger and more sustained beta suppression even before stimulus onset. Notably, this enhancement extended beyond informative cues, suggesting that older adults operate in a chronically elevated state of motor readiness. However, when predictive cues are available, this pre-activated motor system may directly translate prior expectations into action plans, facilitating decision-making but at the expense of sensory evidence evaluation.

**Figure 4.**
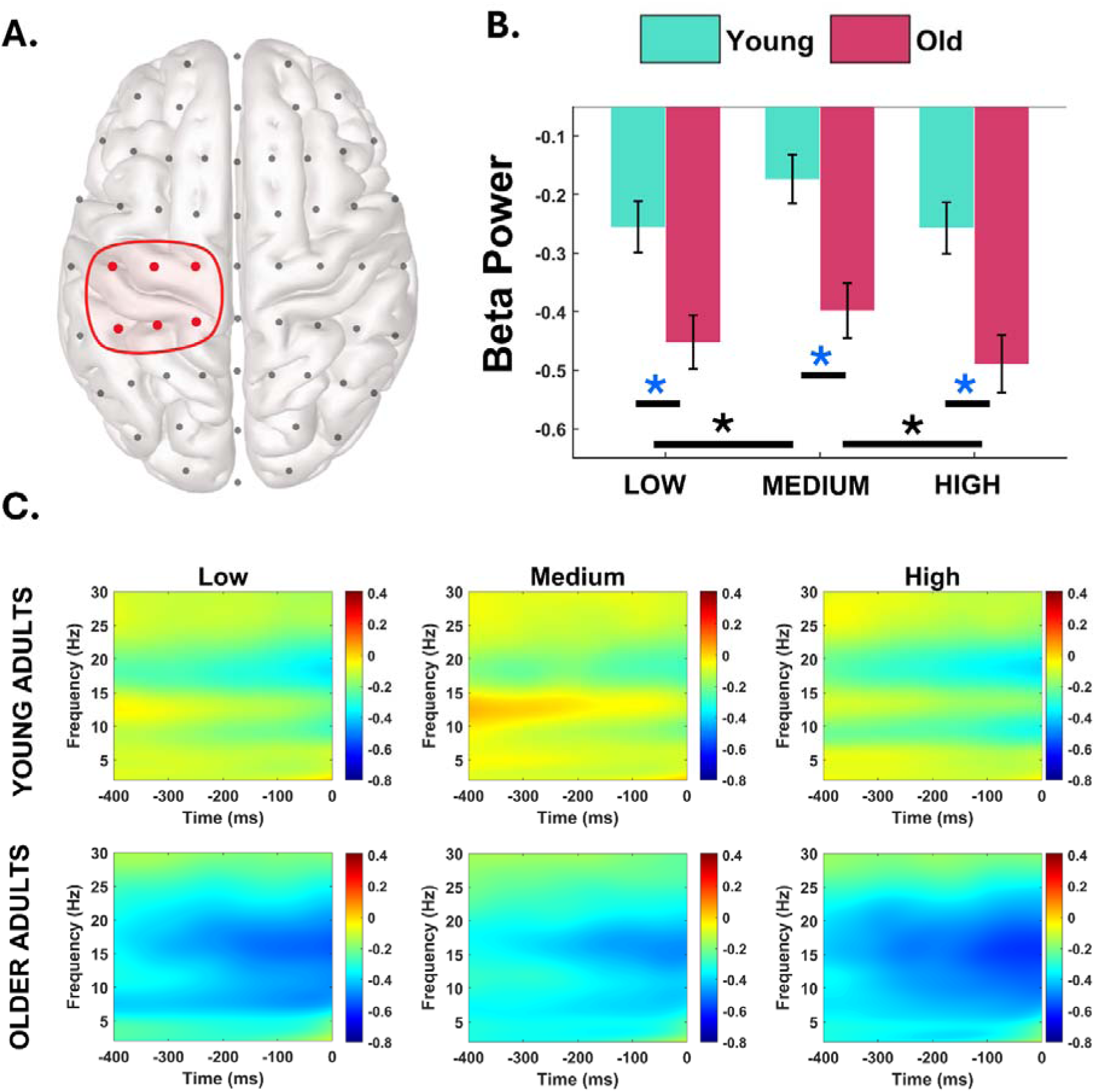
Motor beta___band desynchronization indexes reliance on anticipatory motor strategies in aging. A) Pre-stimulus beta amplitude (14–20 Hz) was extracted left-central electrodes (C5, C3, C1, CP5, CP3, CP1), reflecting contralateral motor cortex engagement for right-hand responses, in low (33 %), medium (50 %), and high (67 %) cue-probability conditions, separately for young and older adults. B) A two-way mixed ANOVA revealed significant main effects of Cue Probability (F_₂_,_₃₀₀_ = 17.32, p < 0.001) and Group (F_₁_,_₁₅₀_ = 12.62, p < 0.001), but no Group × Cue interaction (F_₂_,_₃₀₀_ = 0.70, p = 0.498). Across groups, beta desynchronization was stronger in the informative conditions (black asterisks, high = –0.37 ± 0.04; low = – 0.35 ± 0.04) compared with the medium condition (–0.29 ± 0.04; all t_₁₅₁_ > 4.07, all p < 0.001), with no difference between the two informative cues (t_₁₅₁_ = 1.30, p = 0.196). Older adults exhibited overall greater beta suppression than young participants (blue asterisks, young = –0.20 ± 0.03; old = –0.44 ± 0.03; F_₁_,_₁₅₀_ = 12.62, p < 0.001). An exploratory within-group comparison suggested a trend toward stronger beta desynchronization for high- relative to low-probability cues in older adults (t_₇₁_ = –1.78, p = 0.080). C) Time–frequency representation of pre-stimulus activity (–400 to 0 ms) over left central electrodes. Overall, prior information enhanced motor preparation, with older adults showing a generalized increase in beta suppression, particularly pronounced in the high-probability condition.

### Differential theta engagement in young and older adults during expectation-guided decisions

While sensory and motor oscillations reflect the implementation of expectations at different stages of the decision process, effective perceptual inference also requires executive mechanisms that regulate the influence of prior information. To assess this regulatory component, we examined fronto-parietal theta connectivity, a neural correlate of top-down control^22^, using the weighted Phase Lag Index (wPLI)^24^. We quantified theta-band fronto-parietal connectivity over the same right-hemisphere circuit [frontal ROI (FC2, FC4, FC6, FT8, F2, F4, F6, F8), parietal ROI, (Pz, P2, P4, P6, P8, CP2, CP4, CP6)] identified in our previous work^11^, capturing top-down executive regulation of posterior sensory areas. A two-way mixed ANOVA revealed a significant interaction between group and cue condition (F2,300 = 3.20, p = 0.042), indicating distinct cue-related connectivity patterns in fronto-parietal theta across age groups. Specifically, young adults exhibited stronger fronto-parietal theta connectivity than older adults under both medium- (young = 0.140 ± 0.004, old = 0.126 ± 0.005) and high-probability cues (young = 0.141 ± 0.004, old = 0.128 ± 0.004; all t₁₅₀ > 2.04, p < 0.043), whereas no group difference emerged in the low-probability condition (young = 0.135 ± 0.004, old = 0.129 ± 0.005; t₁₅₀ = 0.91, p = 0.36). Within groups comparisons demonstrated significantly greater theta synchronization in the high-compared to the low-probability condition in young adults (t_79_ = -2.52, p = 0.014). No differences were observed between the two informative cues (all t₇₉ < 1.96, p > 0.054). In contrast, older adults showed no reliable modulation of theta connectivity across cue probabilities (all t₇₁ < 1.06, p > 0.29), suggesting a reduced ability to dynamically adjust executive coupling in response to prior information. To complement the parametric results, we applied a data-driven connectivity index approach drawn by Alekseichuk et al.,^25^. This analysis revealed a significant modulation of theta-band connectivity as a function of prior probability in the young group, with a higher connectivity index observed in the high-probability condition compared to the low-probability condition. Specifically, the proportion of significantly modulated electrode pairs exceeded the 95th percentile of the null distribution derived from 1000 permutations (p = 0.02). In contrast, the elderly group showed no significant difference in connectivity between conditions, with the observed connectivity index falling within the bounds of the permutation-based null distribution (p = 0.81).

**Figure 5.**
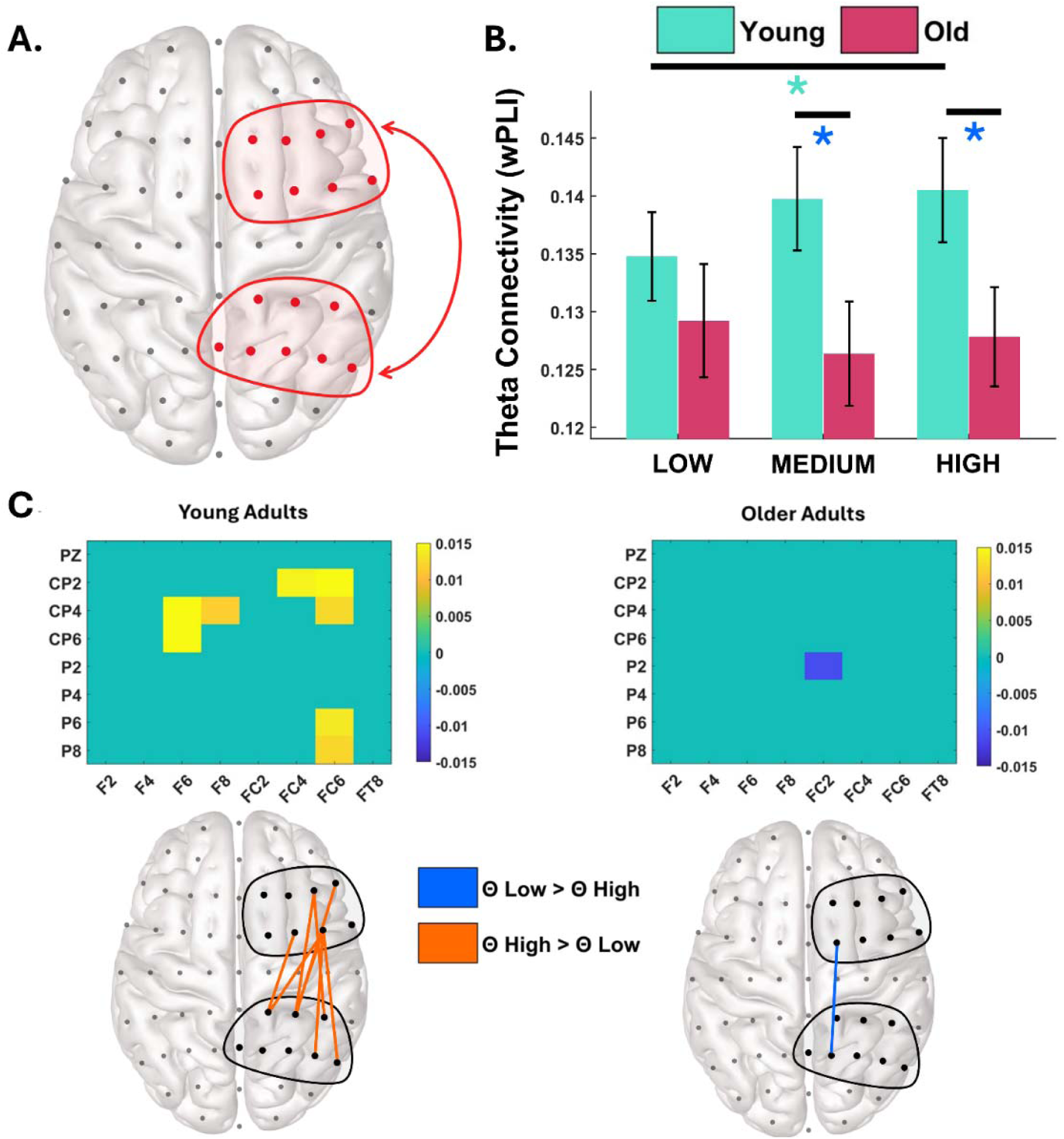
Fronto-parietal theta connectivity during expectation-guided decisions. A) Theta-band fronto-parietal connectivity across cue-probability conditions (low, medium, high) for young and older adults. Connectivity was quantified using the weighted Phase Lag Index (wPLI) between right frontal (FC2, FC4, FC6, FT8, F2, F4, F6, F8) and parietal (Pz, P2, P4, P6, P8, CP2, CP4, CP6) electrodes, reflecting the executive-sensory control pathway identified in prior work. B) A two-way mixed ANOVA revealed a Group × Cue interaction, reflecting age-dependent cue modulation. Between-group contrasts showed stronger theta connectivity in young at medium (blue asterisk, young = 0.140 ± 0.004; older = 0.126 ± 0.005) and high-probability cues (blue asterisk, young = 0.141 ± 0.004; older = 0.128 ± 0.004; all t_₁₅₀_ > 2.04, all p < 0.043), with no difference at low probability (young = 0.135 ± 0.004; older = 0.129 ± 0.005; t_₁₅₀_ = 0.91, p = 0.363). Importantly, young adults showed greater theta connectivity for high vs low probability (green asterisk, t_₇₉_ = 2.52, p = 0.014). Older adults showed no reliable cue-related modulation (all t_₇₁_ < 1.06, all p > 0.292). C) Data-driven connectivity index (proportion of significantly modulated fronto-parietal pairs) computed via a permutation-based approach (1000 iterations), following the approach described in^25^. Theta connectivity was quantified using the weighted Phase Lag Index (wPLI). The young group exceeded the 95th-percentile null threshold for high > low cue probability (p = 0.02), whereas the older group did not (p = 0.81). This data-driven analysis confirms that prior-based modulation of fronto-parietal coupling emerges selectively in the young cohort.

### Age⍰related reorganization of neural dynamics underlying prior⍰driven decision biases

To investigate how the four key oscillatory metrics highlighted - posterior alpha amplitude modulation (high vs. low probability trials), individual alpha frequency (IAF), motor beta desynchronization (informative vs. non informative trials), and fronto-parietal theta connectivity shift (high vs. low probability trials) - jointly underpin the neural implementation of prior-driven decision shifts, we performed a principal component analysis (PCA) separately in each age cohort. This data-driven approach reduces the dimensionality of multiple oscillatory predictors into a small set of latent factors, thereby enabling direct mapping to behavioural indices.

In young adults, the PCA revealed the presence of two principal components. The first (PC1), explaining 29% of variance, captured an antagonistic sensory-executive axis, loading positively on posterior alpha modulation (0.79) and negatively on fronto-parietal theta connectivity (- 0.72). High PC1 scores thus reflected strong prior-related alpha amplitude modulation combined with reduced theta shift between high and low probability conditions, whereas low scores indexed the opposite pattern. Critically, interindividual variability in PC1 positively predicted the magnitude of criterion shifts (ρ = 0.37, p < 0.01). This suggest that the balance between alpha-mediated sensory preparation (which promotes bias) and theta-based executive oversight (which counteracts it) governs the extent to which prior bias decisions in young adults. The second component (PC2), capturing 26% of variance, was dominated by IAF (- 0.83) along with a contribution of beta desynchronization (+ 0.54). While this component did not significantly predict behaviour (ρ = 0.18, p = 0.11), the presence of beta synchronization within this dimension indicates that slower alpha rhythms tend to co-occur with stronger prior-driven motor engagement, even though this association remains functionally secondary in youth.

In older adults, the component structure was reorganized. The first component (PC1, 31% of variance) contrasted posterior alpha modulation (0.78) and theta connectivity (- 0.63), resembling the sensory-executive axis observed in young adults, whereby enhanced alpha modulation was associated with reduced theta coupling. However, this sensory-executive interplay no longer predicted criterion shifts (ρ = 0.10, p = 0.41), suggesting that in older adults the coordination between these rhythms becomes functionally decoupled from behavioural bias. By contrast, the second component (PC2, 26% of variance) was dominated by the prior-driven motor beta desynchronization (0.81), with weaker loadings of theta connectivity (0.43) and IAF (- 0.40). Crucially, PC2 scores significantly predicted criterion shifts (ρ = 0.24, p = 0.04), showing that in aging, prior-driven motor beta desynchronization becomes the main physiological correlate of expectation-driven bias. Notably, the moderate negative loading of IAF also suggests that individuals with reduced intrinsic alpha rhythms exhibited stronger beta suppression and greater behavioural bias, implying that a slowing of intrinsic rhythmicity may bias the system toward a more basic, action-oriented mode of predictive processing, in which expectations are translated into motor preparation rather than proactive sensory analysis of the incoming stimuli. Taken together, these findings reveal a fundamental reorganization of the oscillatory architecture supporting the use of prior information with age. In young adults, decision biases emerge from a distributed mechanism balancing anticipatory sensory processing (alpha amplitude modulation) and executive oversight (theta connectivity), consistent with previous accounts of predictive regulation^11^. In contrast, aging collapses this multi-level coordination into a more unidimensional, motor-centred mode, in which probabilistic information is translated directly into preparatory beta activity. This shift indicates that the ability to exploit prior expectations is preserved in older adults but implemented through a less flexible, more downstream neural route that favour action readiness over proactive sensory tuning.

**Figure 6.**
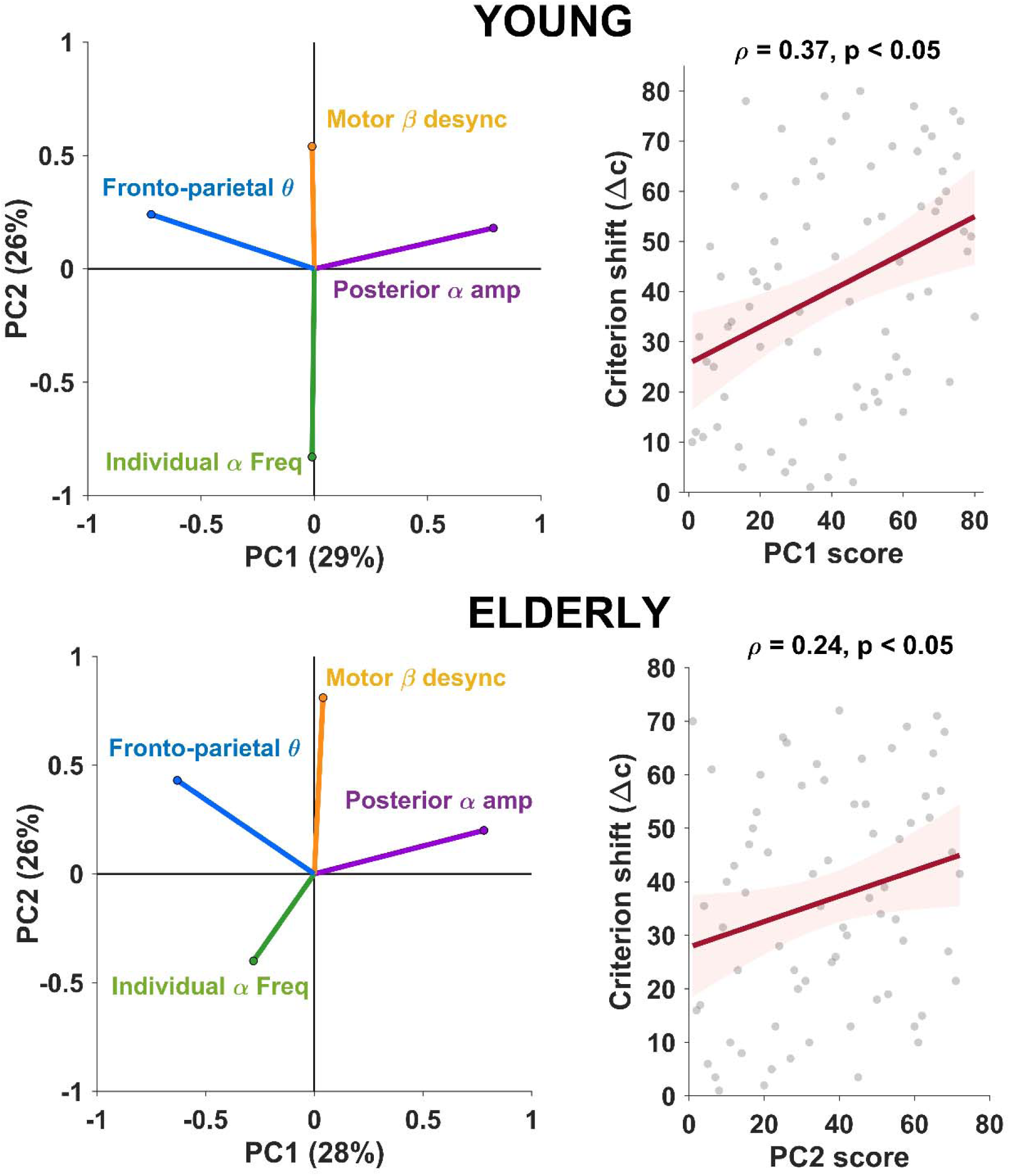
Age-related reorganization of oscillatory components supporting expectation-guided perception. A. (Left Panel) In young adults, PC1 (29% of variance) opposes posterior alpha amplitude shift (purple) and fronto-parietal theta connectivity shift (blue), suggesting the presence of a sensory-monitoring axis. PC2 (26%) instead groups individual alpha frequency (green) and motor beta desynchronization in informative condition (orange), reflecting a secondary motor-IAF dimension independent from PC1 and unrelated to behavioural performance (ρ = 0.18, p = 0.11). (Right Panel) Sensory-monitoring component positively correlates with behavioural bias (ρ = 0.37, p < 0.01), with a weaker theta synchronization combined with a higher alpha modulation connected to increased reliance on prior information in perceptual decision-making. B. (Left panel): In older adults, the latent structure reorganizes into two distinct axes. PC1 (31%) mirrors the sensory-executive configuration observed in young adults, coupling stronger posterior alpha modulation (purple) with reduced fronto-parietal theta connectivity (blue), yet without behavioural relevance (ρ = 0.10, p = 0.41). PC2 (26%), instead, is dominated by motor beta desynchronization (orange), accompanied by a weaker positive theta contribution (blue) and a negative loading of IAF (green). Right panel: PC2 scores correlate with behavioural bias (ρ = 0.24, p = 0.04), indicating that individuals exhibiting stronger prior-driven beta desynchronization, particularly when combined with slower intrinsic alpha rhythms, show greater reliance on prior expectations during perceptual decision-making. This axis captures a shift toward a motor-centred mechanism in which prior information primarily manifests through enhanced beta suppression.

## Discussion

The present study demonstrates that aging profoundly reshapes the neurocognitive architecture supporting expectation-guided perceptual decisions. At the behavioural level, older adults showed a marked shift toward greater reliance on prior information, expressed as increased decision bias (Δ criterion) and reduced perceptual sensitivity (d′) under high-probability conditions. This pattern converges with evidence that aging amplifies dependence on internal predictive models, potentially reflecting a compensatory response to reduced efficiency in sensory evidence processing ^26,27^.

The neurophysiological results parallel this behavioural pattern, revealing age-specific alterations across multiple oscillatory mechanisms that underpin expectation-based perception. These signatures point to a functional reorganization in the coordination among sensory, executive, and motor systems, delineating how aging reshapes the large-scale neural dynamics that support perceptual inference.

First, we observed a clear dissociation in posterior alpha dynamics between age groups. In younger adults, alpha amplitude was modulated by cue probability, with high-probability cues eliciting marked desynchronization relative to low-probability conditions. This pattern reflects the role of alpha oscillations in regulating visual cortical excitability: lower alpha power increases neural excitability, shifting sensory baselines activity and thereby increasing the likelihood of reporting a target ^28^. In high-probability trials, such anticipatory desynchronization biases perception toward expected events, effectively pre-tuning the sensory system for likely stimuli. In contrast, older adults showed no reliable alpha modulation, suggesting a reduced capacity to flexibly adjust visual cortex activity in anticipation of forthcoming stimuli. This result converges with previous findings from spatial cueing paradigms, where older adults - despite successfully allocating spatial attention - failed to exhibit the typical alpha lateralization observed in younger individuals ^29,30^.

Beyond amplitude effects, we observed a robust age-related difference in prestimulus individual alpha frequency (IAF), consistent with prior reports of reduced intrinsic rhythm speed in aging^14,15,31,32^. The effect was independent of cue probability, with no evidence of cue-related modulation in either group. This pattern indicates that, in this context, IAF acts as a stable, trait-like marker of sensory sampling, without any cue-dependent modulation. The slowing of this intrinsic rhythm - previously associated with reduced temporal precision in sensory processing^13,33–35^, poor visual performance ^16,36–38^ and pathological conditions ^39–41^ - may therefore contribute to the behavioural shift observed in aging by fostering greater reliance on prior expectations as a compensatory response to less efficient sensory sampling. Consistent with this view, Hirst et al.^42^ reported that increased susceptibility to the sound-induced flash illusion in older adults was mediated by low-level sensory acuity. Crucially, Cecere et al.,^12^ demonstrated that lower IAF predicts greater susceptibility to the same illusion. This suggests that age-related degradation of unisensory precision broadens temporal integration windows across modalities.

The absence of cue-driven sensory modulation in older adults, despite their pronounced behavioural bias they manifested, suggests a shift in how expectations are implemented: away from anticipatory tuning of sensory cortices and toward downstream motor systems. Supporting this hypothesis, beta dynamics revealed preserved but reconfigured engagement of anticipatory motor processes in aging. Both groups exhibited beta-band desynchronization in informative (high and low probability) relative to non-informative (medium probability) conditions, reflecting proactive motor preparation when prior-congruent actions could be pre-programmed^21,43,44^. However, older adults exhibited a robust increase in beta suppression across all conditions, with the strongest effects in high-probability trials. Notably, this enhancement emerged precisely in the context of reduced perceptual sensitivity and maximal response bias, suggesting that exaggerated beta desynchronization reflects an overcommitment to prior-congruent responses, a premature motor implementation of expectations that bypasses full sensory evaluation^45^. Consistent with this interpretation, Chan et al.^26^ reported greater susceptibility to a sound-induced flash illusion in older adults, accompanied by increased pre-stimulus beta activity compared with younger participants. Within a Bayesian framework, this finding supports the notion that aging is associated with an enhanced reliance on perceptual priors, and that beta-band dynamics may index the neural implementation of these predictions. However, the functional value of this shift may depend on environmental contingencies. In a recent study^18^, stronger beta desynchronization was associated with better performance in individuals who overweight prior information but only in a fully deterministic context. This suggests that older adults, by strategically leaning more on prior-congruent motor activation, might even outperform younger adults in environments where prior information is fully stable and predictive.

Finally, we demonstrated that fronto-parietal theta connectivity, an index of proactive executive control^46,47^, differentiated the two groups in terms of both magnitude and functional flexibility. In younger adults, enhanced theta synchronization in high-probability trials indicated active regulation of prior influence, consistent with the notion of a supervisory mechanism for adaptive bias deployment^11^. These trials likely represent a critical point in the decisional space, where strong priors exert maximal influence and can pull behaviour away from stimulus-driven evidence. In such contexts, it can be hypothesized that additional executive control is required to counteract excessive liberalization of the decision criterion, which increases the risk of false alarms^48^. Notably, this form of regulation has been linked to frontal lobe function, with frontal damage associated with a shift toward overly liberal decision strategies^49,50^. In contrast, in older adults, this regulatory engagement was no longer evident, as theta activity failed to differentiate between cue probabilities, pointing to a decline in the executive control mechanisms that normally constrain the influence of prior expectations.

Building on these oscillatory findings, we applied a Principal Component Analysis (PCA) to capture the dominant axes through which sensory (alpha), motor (beta), and executive (theta) oscillations interact. This approach allows us to move beyond single-band analyses and to characterize how aging reshapes the coordination among these rhythmic systems, revealing distinct large-scale strategies for implementing prior information in the task during perceptual decision-making. In young adults, the PCA revealed an antagonistic alpha-theta axis (PC1), in which stronger prior-driven alpha amplitude modulation (reflecting enhanced sensory preparation) was coupled with weaker fronto-parietal theta regulation (reflecting reduced top-down monitoring on prior integration). This axis was found to be behaviourally relevant in the young group: participants showing a stronger alpha-theta trade-off (i.e., greater alpha suppression and weaker theta connectivity modulation between high- and low-probability cues) exhibited larger shifts in decision criterion, reflecting a stronger reliance on prior expectations. Conversely, participants showing higher theta coupling and correspondingly weaker alpha suppression displayed smaller criterion shifts, consistent with increased executive control over sensory preparation. A second, orthogonal component (PC2) captured a motor preparation axis, characterized by stronger beta desynchronization in informative vs. uninformative trials and lower individual alpha IAF. However, only the alpha-theta trade-off component reliably predicted the extent to which each young participant shifted their decision criterion, indicating that coordinated modulation of sensory (alpha) and executive (theta) rhythms forms the core mechanism supporting expectation-guided perceptual inference in younger adults. In contrast, older adults exhibited a markedly reorganized latent structure. The first component (PC1) preserved the sensory-executive configuration observed in youth i.e., opposing posterior alpha modulation and fronto-parietal theta connectivity. However, this coupling no longer predicted behavioural performance, suggesting that the sensory-monitoring loop remains structurally present but functionally disengaged. The second component (PC2), instead, revealed a motor-centred axis dominated by beta desynchronization, accompanied by a weaker positive contribution of theta and a negative loading of individual alpha frequency. This component uniquely predicted criterion shifts in older adults, indicating that stronger prior-driven beta suppression underlies greater behavioural bias. Notably, the negative contribution of IAF suggests that individuals with lower intrinsic rhythms, reflecting poorer sensory sampling^13,16^, exhibited stronger beta modulation. This reorganization points to a functional reallocation of predictive processing with age. In aging, predictive processing appears to shift toward the motor domain, where beta activity provides the primary channel for expressing prior information. The concurrent reduction of IAF aligns with this reconfiguration: as neural rhythms supporting fine-grained sensory sampling deteriorate, the system increasingly relies on lower-level, motor-based mechanisms that remain efficient and accessible.

Together, these results reveal a functional reorganization of predictive mechanisms with age. In young adults, predictive inference arises from a distributed coordination between sensory (alpha), executive (theta), and motor (beta) rhythms, a flexible network in which alpha suppression enables anticipatory sensory sampling, theta provides adaptive control, and beta supports preparatory motor readiness. In older adults, this distributed structure collapses into a more compact configuration: alpha and theta dynamics disengage from the predictive loop and beta desynchronization emerges as the dominant carrier of prior implementation. In this framework, beta activity reflects a form of preparatory predictiveness: an automated readiness state that replaces flexible sensory preparation/monitoring with a more direct, embodied implementation of prior information.

From a predictive coding perspective, this reorganization may reflect an imbalance in the hierarchical estimation of precision - that is, in the inferred reliability of top-down versus bottom-up signals^51,52^. With age, this balance appears to shift toward increased precision-weighting of internal models and reduced weighting of sensory evidence, consistent with the weakening of alpha-based mechanisms. The accompanying dominance of beta activity suggests that prior expectations are no longer flexibly tested against incoming input but are instead implemented as anticipatory motor templates. Such a redistribution of hierarchical precision would render perceptual inference more rigid and self-reinforcing, predisposing older observers to rely on habitual, expectation-driven prediction rather than dynamically updating beliefs in response to sensory uncertainty. Interestingly, this imbalance between internal prediction and sensory updating resembles patterns observed in clinical conditions characterized by altered perceptual inference^10,53–55^, although in aging it likely reflects an adaptive, compensatory reorganization rather than a pathological deviation.

Notably, this framework highlights a shift not merely toward decline but toward a compensatory regime in which neural resources are strategically reallocated to sustain performance despite sensory degradation. Yet, as resource constraints intensify, these compensatory processes may lose flexibility and become structurally rigid, ultimately increasing vulnerability to cognitive inefficiency. From a theoretical standpoint, the present findings refine current models of cognitive aging by delineating the oscillatory mechanisms through which prior expectations are integrated, and progressively over-weighted, in older adults. Rather than signalling maladaptation, the behavioural bias observed here may represent a trade-off in which efficiency and predictability are prioritized over accuracy when sensory evidence becomes unreliable. Notably, chronological age within the older group was unrelated to behavioural or neural measures, indicating that the observed variability reflects qualitative differences in predictive strategy rather than progressive deterioration.

In conclusion, our study shows that aging reshapes perceptual decision-making via a functional reorganization of the neural mechanisms supporting the integration of sensory evidence and prior expectations. By characterizing age-related changes across sensory, motor, and executive domains, we provide a mechanistic account of how older adults adapt their decision strategies, shifting toward a more anticipatory, prior-driven mode of processing. These findings highlight the brain’s capacity to adaptively restructure decision-making strategies in later life, shifting from flexible sensory-executive dynamics to more streamlined, prior-based routes. Rather than reflecting decline, this transition could represent a strategic reconfiguration tailored to the constraints of the aging system, optimizing decisional efficiency under ambiguous conditions.

## Methods

### Participants

Participants consisted of two groups: a group of young adults (N = 80; mean age = 23.78, SD = 2.79; 43 females) and a group of older adults (N = 72; mean age = 64.93, SD = 7.42 years; 33 females). Participants were rigorously screened to exclude neurological, psychiatric, or sensory disorders, and provided written informed consent following ethical guidelines approved by the local ethics committee (protocol code 201723, approved on 26 August 2021), consistent with the Declaration of Helsinki.

### Task and Procedure

Participants performed a probabilistic detection task specifically designed to investigate perceptual decision-making under varying degrees of sensory expectations. The experimental design consisted of two distinct phases. In the first phase, each participant underwent an adaptive psychophysical titration procedure aimed at determining the individual contrast threshold for detecting grey circular stimuli embedded within each square of a black and white checkerboard. This procedure ensured approximately 70% detection accuracy under conditions with equal numbers of target-present and target-absent (catch) trials ^38^. Following the titration phase, participants completed six experimental blocks, each containing 90 trials. Each trial began with a visual probabilistic cue, signalling the likelihood (low, medium, or high) of the subsequent appearance of the target stimulus (grey circles embedded within a checkerboard pattern) located in the lower-left part of the screen. After a random interval between 1.2 and 1.5 seconds from cue offset, the checkerboard appeared, either containing the target stimuli or remaining empty (catch trial). Participants indicated the presence (pressing the "K" key) or absence (pressing the "M" key) of the target using their right hand, with no time constraints to avoid speed-accuracy trade-offs. After the participant’s response, an inter-trial interval with a random duration between 1.9 and 2.4 seconds occurred (gray screen). The probabilistic cue was visually represented by a rectangle partially shaded in red (bottom portion) and partially in blue (top portion). The proportion of red shading indicates the probability of target occurrence. Specifically, the cue conditions included: high probability (67% likelihood), low probability (33% likelihood), and medium probability (50% likelihood, representing no predictive information). The actual frequency of stimulus presentations corresponded exactly to cue probabilities, a factor explicitly communicated to participants.

### Behavioral Data Analysis

Behavioral data were analysed using Signal Detection Theory (SDT) ^56^ to calculate perceptual sensitivity (d’) and response criterion (c). The sensitivity measure (d’) quantifies participants’ ability to discriminate between target-present and target-absent trials, with higher values indicating greater perceptual accuracy. The criterion measure (c) quantifies response bias, indicating a tendency to favour either affirmative (liberal) or negative (conservative) responses; values significantly different from zero suggest biased decision-making. Both indices were computed from hit rates and false alarm rates. To assess whether sensitivity and criterion were modulated by cue probability, and whether such modulation differed between age groups, statistical analyses employed separate two-way ANOVAs with Group (2 levels: young, older adults) and Cue probability (3 levels: low, medium, high) as fact. Significant interactions were further explored using post-hoc comparisons. In addition, to index the cue-induced bias shift, we computed Δcriterion = c_low − c_high for each participant and compared Δ criterion between groups using a two-sample (independent) t-test.

### EEG Data Acquisition and Preprocessing

EEG data were recorded using a 64-channel cap arranged according to the international 10–10 system at a sampling rate of 1000 Hz. Offline preprocessing included band-pass filtering (0.5–100 Hz) and notch filtering (49–51 Hz) to remove line noise, followed by interpolation of noisy electrodes and visual rejection of epochs containing residual artifacts. Ocular components were identified and removed using Independent Component Analysis (ICA). The data were then re-referenced to the average reference, downsampled to 256 Hz, and subjected to a surface Laplacian transform (spherical splines) to enhance spatial specificity.

### EEG Analysis

Time–frequency decomposition was performed using complex Morlet wavelet convolution applied to the continuous EEG time series, extracting time-resolved amplitude estimates at each target frequency. Each wavelet was defined as a complex sine function tapered by a Gaussian envelope. A family of wavelets was generated for frequencies between 2 and 30 Hz, with the number of cycles increasing from three (lowest frequencies) to eleven (highest frequencies), thereby balancing temporal and spectral resolution. EEG data and wavelets were transformed into the frequency domain using Fast Fourier Transform (FFT), multiplied in the frequency domain, and then converted back to the time domain via inverse FFT. Amplitude was obtained as the absolute value of the resulting complex signal, while phase was defined as the angle of the complex output. Amplitude values were baseline-corrected using decibel (dB) conversion relative to the mean amplitude in the –3100 to –2700 ms window preceding stimulus onset.

### EEG Oscillatory Analysis

The EEG analyses focused on alpha, beta, and theta oscillations, guided by prior evidence demonstrating their specific roles in perceptual expectation integration.

#### Alpha amplitude analysis

To quantify preparatory activity in visual sensory areas, alpha amplitude (7–13 Hz) was extracted in the −400 to 0 ms pre-stimulus window over right-posterior electrodes (Oz, POz, Pz, O2, PO4, PO8, P2, P4, P6, P8), following the approach described in^9–11^. This lateralized ROI was selected a priori based on (i) its maximal alpha reactivity in our previous work^9^ and (ii) the fact that the target stimulus was always presented in the left visual hemifield, thereby predominantly engaging right-hemisphere visual areas. A two-way mixed ANOVA was conducted with Group (young, older adults) as the between-subjects factor and Cue Probability (low, medium, high) as the within-subjects factor to examine age-related differences in pre-stimulus alpha power across varying levels of cue probability. Post-hoc analyses were performed to explore significant interactions.

#### Individual Alpha Frequency analysis

We estimated pre-stimulus IAF separately for each cue condition in the −400 to 0 ms window. To ensure consistency with the alpha amplitude analysis, IAF was estimated over the same right posterior ROI contralateral to the stimulus location (Oz, POz, Pz, O2, PO4, PO8, P2, P4, P6, P8). For each electrode and condition, we computed power spectra with EEGLAB’s *spectopo* function and removed the aperiodic 1/f background by fitting a log–log linear model of power. The periodic-only spectrum (in dB) was obtained by subtracting the fitted aperiodic component. After applying a *Savitzky–Golay* smoothing, the alpha peak was automatically searched within 7–13 Hz for each channel. Edge peaks (exactly at 7 or 13 Hz) were flagged and excluded. The IAF for a given participant and condition was taken from the posterior electrode with the highest mean alpha power that yielded a non-edge alpha peak. All automatically detected peaks were visualized and implausible estimates (e.g., residual edge peaks) were corrected by visual inspection. The resulting IAF values entered a two-way mixed ANOVA with Group (young, older adults) and Cue Probability (low, medium, high) as factors.

#### Beta amplitude analysis

Motor beta-band activity (14–20 Hz), a well-established marker of motor preparatory activity^21^, was quantified in the pre-stimulus window (–400 to 0 ms)^18^ over a left central region of interest (C5, C3, C1, CP5, CP3, CP1), corresponding to the hemisphere contralateral to the response hand. This ROI was selected a priori based on the well-established lateralization of beta desynchronization over motor cortex during right-hand actions^18,19,57,58^. To test how prior expectations modulate motor preparation between the two groups, a two-way mixed ANOVA was conducted with Group (young, older adults) as the between-subjects factor and Cue Probability (low, medium, high) as the within-subjects factor. This analysis tested for beta amplitude differences across the three levels of cue probability (low, medium, high), allowing us to assess whether beta desynchronization scaled with the informativeness of the cue and whether this pattern differed between groups.

#### Theta Connectivity Analysis

Fronto-parietal theta-band connectivity, associated with executive control mechanisms, was quantified using the weighted Phase Lag Index (wPLI) between the right frontal (FC2, FC4, FC6, FT8, F2, F4, F6, F8) and parietal (Pz, P2, P4, P6, P8, CP2, CP4, CP6) electrode pairs in the pre-stimulus window (–400 to 0 ms) following the approach employed in Tarasi et al.,^11^ and replicated in Sunder et al.^59^. This lateralized fronto-parietal network was selected a priori based on our working hypothesis that executive regions modulate posterior sensory areas engaged by the task (i.e., the right hemisphere as sensory processing of the left visual field (where stimuli were presented) predominantly engages right posterior cortex), and on previous evidence from our earlier study showing expectation-driven theta coupling in the same right-lateralized circuit^11^. Specifically, in Tarasi et al.^11^, theta synchronization was shown to increase in high-probability compared to low-probability conditions, and this modulation predicted a controlled integration of prior information during perceptual decision-making. The wPLI ^24^ is an advanced version of the Phase Lag Index (PLI), which quantifies connectivity as the absolute value of the average sign of phase angle differences (+1 or −1 relative to the real axis). Unlike PLI, wPLI places greater weight on phase differences far from the real axis and thus mitigates the influence of spurious synchrony due to volume conduction. wPLI values range from 0 (random phase relationship) to 1 (perfect phase synchronization).

To assess whether prior expectations modulated pre-stimulus brain connectivity in the fronto-parietal network, a two-way ANOVA was conducted with Group (young, older adults) as the between-subjects factor and Cue Probability (low, medium, high) as the within-subjects factor. Additionally, to complement the parametric analysis, we applied a data-driven procedure inspired by Alekseichuk et al., ^25^. For each pair of electrodes within the regions of interest, we conducted a paired t-test comparing wPLI values between the high- and low-probability conditions. A connectivity index was then computed as:

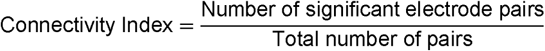

To evaluate statistical significance, a non-parametric permutation test was conducted. For each participant, condition labels (high vs. low probability) were randomly permuted, and the connectivity index was recalculated on the shuffled data. This procedure was repeated 1000 times, generating a null distribution of *Connectivity Index* against which the real one was compared. Connectivity indices exceeding the 95% confidence interval of the null distribution were considered statistically significant. This analysis was conducted separately for the young and elderly groups.

Finally, we examined whether neural and behavioral indices covaried with chronological age within the older cohort. No significant correlations were observed (all *p* > 0.17), indicating that within-group variability was not merely explained by chronological age.

### Principal Component Analysis

To capture the principal modes through which expectation-sensitive oscillatory signals converge into coherent neural strategies, we applied an exploratory PCA to four neural metrics extracted, performing the analysis separately for the young and older cohorts. For each rhythm, we selected the metric that best captured its functional role in prior use. Specifically, we computed (1) posterior alpha modulation (difference in pre-stimulus alpha amplitude between high and low probability cues), (2) motor beta shift (contrast in beta desynchronization between informative [mean(high, low)] and medium cues), (3) fronto-parietal theta shift (difference in theta connectivity between high and low probability cues), and (4) individual alpha frequency (IAF) as a measure of intrinsic sampling rate. For alpha, beta, and theta, we used cue-induced modulation indices, as these oscillations represent state-dependent mechanisms dynamically adjusting to prior information. In contrast, for IAF we used the mean value across cue conditions, reflecting its trait-like nature as an index of intrinsic temporal resolution rather than a transient task-induced modulation. Indeed, in our dataset, alpha, beta, and theta measures showed clear cue-dependent effects (main or interaction effects involving Cue or Group × Cue), directly indexing the neural implementation of prior-driven processes such as sensory preparation, motor preparation, and executive gating, respectively, whereas IAF exhibited a robust main effect of Group (older < young) with minimal within-subject modulation. Components with eigenvalues > 1 were retained. Individual component scores, computed as the linear combination of original metrics weighted by their rotated loadings, were then correlated with behavioural Δ criterion to determine which latent oscillatory dimensions predicted prior-induced decision biases.

